# Differential Compositional Variation Feature Selection: A Machine Learning Framework with Log Ratios for Compositional Metagenomic Data

**DOI:** 10.1101/2021.12.08.471758

**Authors:** Andrew L. Hinton, Peter J. Mucha

## Abstract

The demand for tight integration of compositional data analysis and machine learning methodologies for predictive modeling in high-dimensional settings has increased dramatically with the increasing availability of metagenomics data. We develop the differential compositional variation machine learning framework (DiCoVarML) with robust multi-level log ratio bio-marker discovery for metagenomic datasets. Our framework makes use of the full set of pairwise log ratios, scoring ratios according to their variation between classes and then selecting out a small subset of log ratios to accurately predict classes. Importantly, DiCoVarML supports a targeted feature selection mode enabling researchers to define the number of predictors used to develop models. We demonstrate the performance of our framework for binary classification tasks using both synthetic and real datasets. Selecting from all pairwise log ratios within the DiCoVarML framework provides greater flexibility that can in demonstrated cases lead to higher accuracy and enhanced biological insight.

## Introduction

Ubiquitous use of rapidly advancing metagenomic sequencing technologies are allowing researchers to uncover profound insights into precisely how alterations of microbial communities that live in and on the human body are associated with human disease. These promising technologies are being rapidly explored for use as non-invasive diagnostic and screening tools.^1,2^. The success of such efforts relies on the identification of robust microbial biomarkers that are predictive of disease onset and/or progression. To this end, researchers are exploring novel ways to apply the latest advances in supervised machine learning methodology to unlock key biomarker signals across unique and costly metagenomic data.^3^ Fortunately, numerous publicly-available curated metagenomic datasets^4–6^ have helped researchers develop and test new methodologies on complex metagenomic datasets across diverse disease groups.

In order to discover generalizable metagenomic biomarkers, new prediction methods will need to address important statistical challenges related to high-dimensional metagenomic data. In particular, whole genome shotgun (WGS) and 16S ribosomal-RNA (16S) sequencing techniques utilized in metagenomic studies produce count data that have arbitrary limits on the total number of reads obtained for each sample by the instrument.^7^ Because of this, analysis of these data is limited to relative rather than absolute comparisons. Data with such constraints, formally known as compositional data,^8^ have a simplex sample space and, notably, important dangers arise when ignoring compositional constraints during analysis, including non-linear distances between samples as subsets of parts (e.g. taxa, metabolites) change, spurious correlations, and generalizability of models.^7^ Additionally, sparsity, discreteness, and distribution of total library sizes can influence conclusions and further complicate analyses.^9^ The importance of appropriately analyzing compositional metagenomic data has been discussed extensively in Refs. 7, 10 and the need for suitable normalization approaches to account for these challenges has been highlighted in Ref. 11.

Supervised classification predictive modeling in metagenomics studies aims to classify disease based on learned patterns in microbial compositions. These models can then be deployed in non-invasive screening, diagnostic, or prognostic tests. It is not uncommon for researchers to train standard machine learning algorithms such as random forests, support vector machines, or LASSO regularized logistic models on untransformed relative abundance data and group labels (e.g. disease, phenotype) to identify microbial compositional differences between groups of interest. Indeed, standard approaches such as those in MetaML^12^ ignore compositional data constraints intrinsic in relative abundance data; as a result, model interpretability and beyond-study generalizability may be severely limited. Fortunately, various log-ratio transformations have been proposed to overcome these challenges.^8,13^ In particular, given signals from *p* parts, one seeks to properly describe the corresponding point on the (*p* − 1)-dimensional simplex. Only the (*p* − 1)-feature basis from the additive log ratio (ALR) transformation and the spanning frame of all 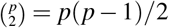 features from the pairwise log ratio (PLR) transformation provide both simple interpretation^14^ and subcompositional coherence.^8^ We note in particular that the ALR transformation has been shown to be an effective way to statistically analyze omics data^15^ and that variable selection approaches that address compositional constraints on metagenomic data were important to the recently proposed selbal^16^ and coda-lasso^17^ methods.

We here introduce the differential compositional variation machine learning framework (DiCoVarML) for guided feature selection to efficiently train, robustly test, and flexibly interpret supervised classification models using robust additive log ratio or pairwise log ratio features. To more fully motivate the utility of this framework, we first demonstrate important generalization limitations of machine learning models that are trained directly with either relative abundance or centered log ratio (CLR) transformed data: in a simple low-dimensional setting, even with proper cross-validation performance estimation, AUC estimates obtained for models trained with relative abundance or CLR transformed data may be unreliable. We then show through detailed simulation study and analysis of publicly-available metagenomic datasets that DiCoVarML provides significantly better classification performance than existing compositional feature selection approaches. To demonstrate its clinical utility, we then apply DiCoVarML to predict the onset of necrotizing enterocolitis in premature infants using fecal metagenomic data from the NICU NEC study,^18^ and we develop a novel meta-analysis covering 9 studies^19–27^ to classify microbial differences between the gut microbiome and colorectal cancer.

## Results

### The DiCoVarML framework

DiCoVarML attempts to obtain an optimal set of log ratios between parts (e.g., taxa, metabolites) for use as features in machine learning classification models. Broadly speaking, feature selection for compositional metagenomic data analysis can be applied at either (a) the parts level (selbal, clr-lasso, coda-lasso) or (b) the log-ratio level (ALR or PLR). In DiCoVarML, we utilize a novel multi-level feature selection approach to simultaneously identify a robust subset of log ratios between a targeted number of parts. Our versatile approach importantly supports both targeted (number of target parts selected by expert) and untargeted (number of selected parts optimized empirically) feature selection. A targeted approach is especially useful when weighing trade-offs between predictive performance and cost for diagnostic test development, where limiting the number of parts to include in the final assay may be of particular concern. Moreover, depending on study priorities (high interpretability, high prediction), the DiCoVarML framework naturally allows for either interpretable but generally less accurate ridge-regression models or complex but generally more accurate average probability ensemble models for classification. Additionally, biomarker signatures discovered by DiCoVarML can be found via ALR (lower computational cost but possibly lower insight) or PLR (higher computational cost for possibly higher insight).

To robustly select features and estimate performance while minimizing overfitting, the DiCoVarML framework uses a double-nested (inner and outer) *k*-fold cross-validation learning schema to discover log-ratio signatures and estimate classification performance (Figure 1a & 1b). That is, the relative metagenomic dataset is randomly split in the outer cross-validation loop into *k* discovery and test partitions with appropriate stratification (by class, sample, study, etc.) given the study design (cross-sectional, longitudinal, repeated measures, etc.) (Figure 1a) with each fold held out and used once as a test set and the remaining folds used for discovery and training. On each partition, the targeted multi-level feature selection (tarMFS) method (Figure 1b) is applied, yielding user-selected classification performance measures (e.g., AUC, accuracy, AUPRC, etc.). To discover an appropriate number of parts (targeted or untargeted) and classification model (from a set selected *a priori* by the user; ensemble and ridge regression are considered here), an inner cross-validation loop is used (Figure 1b): the discovery set of the outer loop is further split into *k*_inner_ folds (here *k*_inner_ = 2) for training and validation. Each classification model (for numbers of parts listed in the array *T*) is evaluated on each partition using the targeted discovery mode (Figure 1c, see “Targeted multi-level feature selection” in Methods). The best performing model and number of parts (*m*_*max*_ and *t*_*max*_ in Figure 1b) is selected and used to train the discovery set model and classification performance is then obtained on the corresponding test fold. Overall classification performance is then estimated by averaging the performance across discovery-test folds (Figure 1a). Depending on study objectives and computational resources, the entire process can be repeated under different random splits to better estimate out-of-study classification performance.

**Figure 1.**
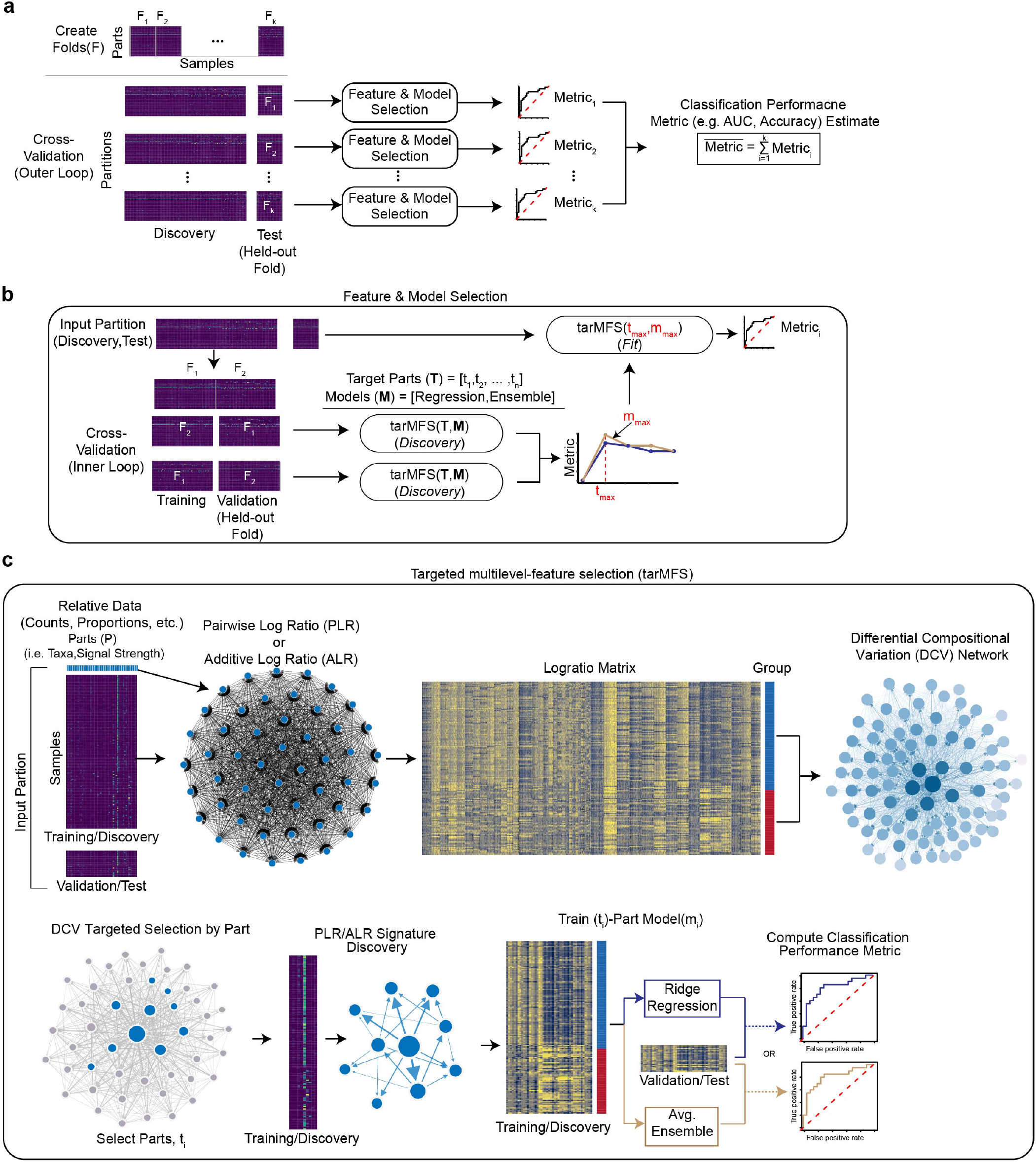
The differential compositional variation machine learning (DiCoVarML) framework for high dimensional compositional metagenomic data. (a) High level overview of the DiCoVarML framework showing the outer cross-validation procedure for estimating out-of-sample performance including the data partitioning, feature/model selection, and classification performance metric estimation. (b) Overview of the partition-specific feature and model selection process showing how targeted multilevel-feature selection (tarMFS) is used in the inner loop to select the number of parts and the model to then be used in the outer loop to estimate classification performance. (c) Overview of the nested targeted-multilevel feature selection method for identifying key parts, log ratio signatures, and estimating classification performance.

### Beyond-study generalization limitations of using relative abundance or CLR-transformed data

Using a simple three-part composition (*p, q, r*) toy dataset, we demonstrate here that machine learning models trained with relative abundance or CLR-transformed count data can easily fail to generalize beyond the available training samples, but that nevertheless the use of additive (ALR) or pairwise log ratios (PLR) succeeds in such settings. In particular, we demonstrate this in scenarios with and without feature selection. We simulate two distinct classes of data across two partitions (train, test) with a common decision boundary within the simplex. While the train and test data partitions share decision boundaries in our simulated data, the deliberate geometric separations between the two simulated partitions are designed to represent differences as might arise from variabilities in study design, sample preparations, different instruments, noise from measurement error, or other cross-study differences. The three-part composition yields a two-dimensional simplex sample space, with the relative proportions of the three parts visualized in the ternary diagrams of Figure 2.

**Figure 2.**
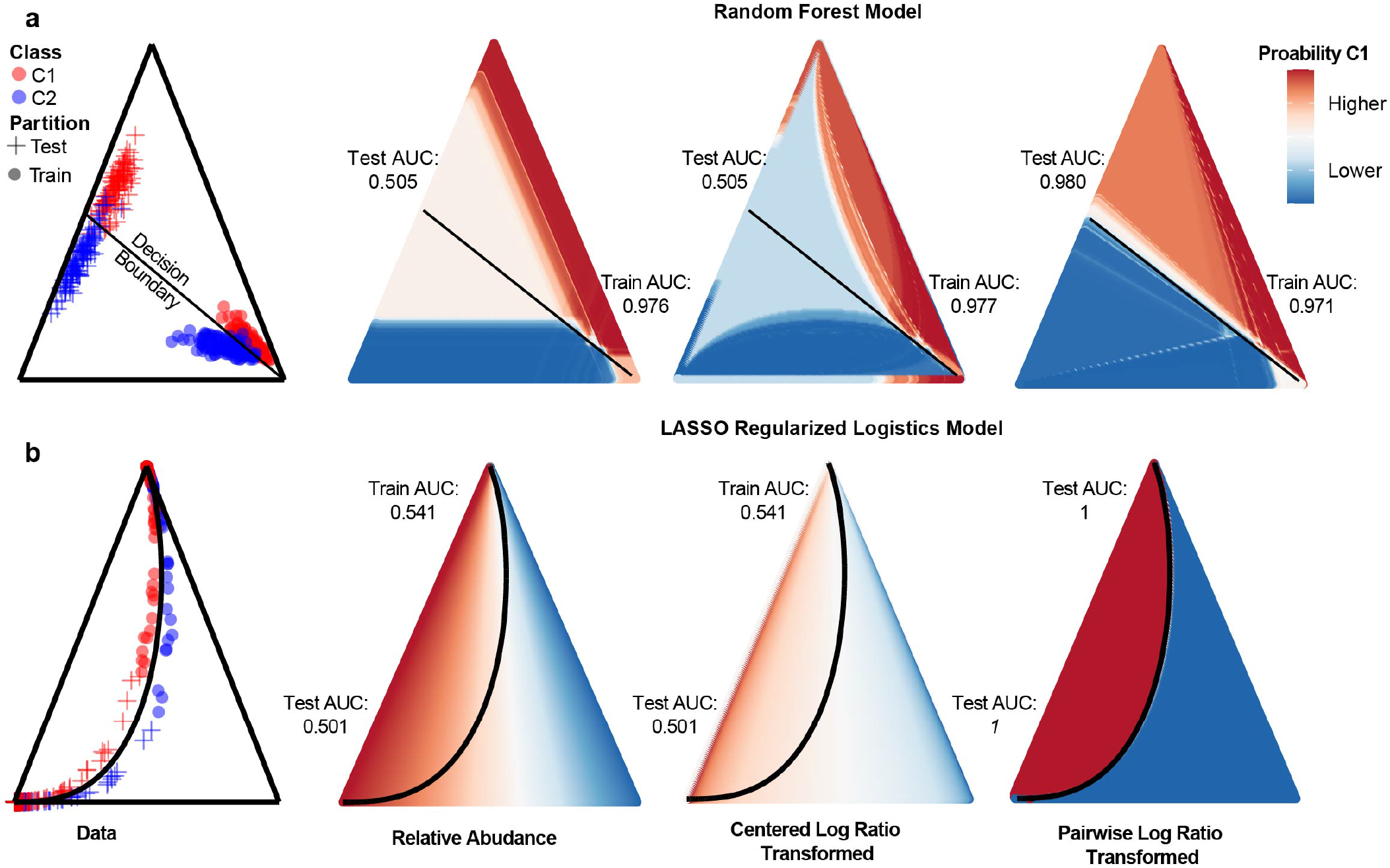
Generalizability limitations from training machine learning models with relative abundance or centered log ratio data. Ternary diagrams visualize compositional data from three component parts in barycentric coordinates. For each scenario (row), data were simulated from two classes (C1, C2) with distinct data distributions for training (circles) and test (+ symbols) partitions in the first column on the left. The corresponding global decision boundaries are shown as solid lines for each scenario. Training set and test set AUC are shown for models trained from relative abundance (second column), centered log ratios (third column), and all pairwise log ratios (right column). Prediction probabilities for class C1 for each model fit to the training partition are indicated by shaded coloring throughout the simplices. (a) Random forest models fit to normally-distributed data on the 2-simplex with an isoproportional decision boundary. (b) LASSO regularized logistic model fit to data separated by an *α* ⊙ {0.6, 0.3, 0.1} compositional line (through barycenter) decision boundary.

We first evaluated a scenario without feature selection where two classes of data are simulated from the additive logistic normal distribution (see Methods) with a single noisy part (Figure 2a). In this scenario, the decision boundary of the (*p, q, r*) components is an isoproportional *p/r* = 1 line in the simplex. Using a random forest model trained directly on the three-part relative abundance (proportions) data, we observed a mean cross-validated AUC = 0.976 in the train set; however, after applying this trained model to the test set, the generalization performance on the test set drops to AUC = 0.505. As observed clearly in the Figure, the random forest model trained on relative abundances failed to learn the correct decision boundary except in the area immediately local to the training set. Similarly, the corresponding model trained on the CLR transformation of the three-part composition yielded a train set AUC = 0.977 but test set AUC = 0.505. In contrast, training the random forest model with all pairwise log ratios (PLR) leads to a train set AUC = 0.971 with test set AUC = 0.980, demonstrating accurate beyond-sample generalization from using pairwise log ratios. In contrast to these stark performance differences, we note that LASSO logistic regression using any of these considered data transformations performs well on this simple, low-dimensional isoproportional boundary scenario.

We next examined a scenario with feature selection on data simulated on opposite sides of a compositional line decision boundary specified by a [0.6, 0.3, 0.1] leading compositional vector through the barycenter (see “Compositional line decision boundary” in Methods). In this scenario, we trained and tested a regularized logistic model on each data transformation to evaluate its generalization properties (Figure 2b). For both relative abundances and CLR transformed counts we obtained train set AUC = 0.541 and test set AUC = 0.501, indicating significant under fitting. That is, in each of these cases it appears that the regularization eliminated the degrees of freedom needed to reconstruct the true log ratios that combine to form the simulated decision boundary. In contrast, when working with the full frame of all pairwise log ratios, the regularized regression achieves (empirical) train set AUC = 1 and test set AUC = 1. We additionally note that all of the considered data transformations generalize poorly when random forest models are trained without feature selection in this scenario, emphasizing the importance of feature selection.

Whereas the use of either pairwise or additive log ratios generalizes well beyond the training sample, we note with these simple illustrative scenarios that there are important beyond-study generalization dangers when using either relative abundances or CLR-transformed data to train machine learning models. Importantly, these limitations are independent of the number of dimensions and can be present (or even expected) in high-dimensional metagenomic data. Notably, the ability of a classification model to generalize true patterns beyond the study immediately at hand is essential both to understanding how parts contribute to a classification and to developing robust diagnostic assays.

### Classification performance evaluation using synthetic data

To understand how untargeted DiCoVarML with PLR, ALR, or hybrid signatures compares to existing compositional approaches with feature selection (selbal, clr-lasso, coda-lasso) we generated synthetic counts mimicking WGS and 16S binary-class data (see Methods) and compared the classification performance of each method (Figure 3). In particular, we studied three scenarios with increasing signal density (number of associated parts) from 2% to 50% of all taxa (WGS = 270, 16S = 124). For each scenario, we increased the percent mean difference (mean shift) between associated taxa from 18% to 30% (see Methods). As seen in Figure 3, for the 16S data with 2% signal sparsity, clr-lasso and selbal performed better than DiCoVarML at the 18% mean shift level, comparably at the 24% level, and slightly worse at the 30% level. As signal density increased, DiCoVarML outperformed the other methods and maintained consistent performance across all signal sparsity settings. Notably, for existing methods on the 16S data, we observed decreasing performance as the signal density increased. These findings are also consistent with simulation results reported in Ref. 17. Notably, coda-lasso performed poorly across all 16S scenarios tested. For the WGS data, similar trends were observed with existing methods (including coda-lasso) performing better or similar to DiCoVarML at the lower signal sparsity levels but worse as the signal density increased. For existing methods on WGS data, we again observed large reductions in performance as signal density increased. Moreover, for DiCoVarML signatures on WGS data, we note a general but smaller reduction performance as signal density increases. Notably, ALR-derived signatures performed worse than PLR and Hybrid signatures in complex WGS data but similarly on less complex 16S data. From this, our simulation results demonstrate DiCoVarML maintains consistent performance over a range of scenarios with 16S or WGS characteristics when compared to existing methods.

**Figure 3.**
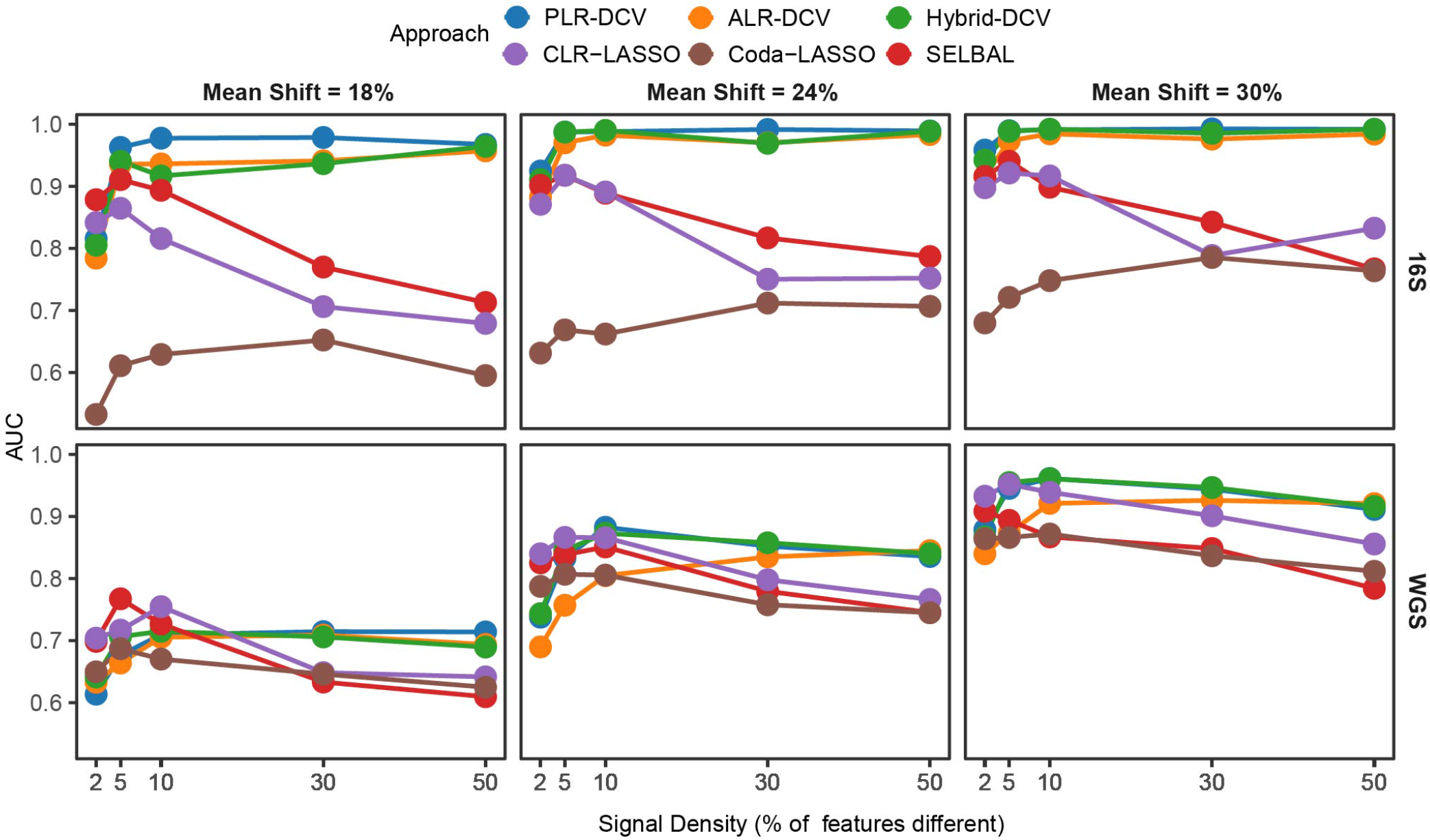
Classification performance comparison using synthetic data distributions. Mean binary classification AUC (5 repeats x 2-fold cross validation) for each method applied to simulated two-class data (n = 100 per class) corresponding to 16S (top row; 165 features) or WGS (bottom; 270 features) characteristics. Signal density (x-axis) measures the percentage of total parts that are different between classes. Columns represent effect size (percentage of between-class mean differences) of each signal feature.

### Classification performance evaluation using real metagenomic data

We next compared the binary classification performance of untargeted DiCoVarML signatures to existing approaches using real case-control 16S and WGS gut microbiome datasets. To better assess the versatility of each approach, we tested classification performance in eight unique disease settings across eight cohorts from four publicly-available data sources (see Methods). Using a paired design, we trained and tested each approach on the same partitions using 15 repeats of 2-fold cross-validation. For ease of interpretation here, we characterize paired mean AUC differences (Δ = AUC_*DiCoVarML*_ − AUC_*existing*_) between approaches as: “modest” ∈ (0.00, 0.01], “moderate” ∈ (0.01, 0.05], and “large” *>* 0.05.

For 16S datasets, DiCoVarML signatures achieved the highest mean AUC in all four datasets tested (Figure 4A) when compared to existing methods. Particularly, ALR signatures achieved the highest AUC on three of four 16S datasets with the Hybrid signature doing best on the remaining dataset. To understand the magnitude and significance of differences in AUC between the existing approaches and DiCoVarML, we used Wilcoxon signed rank tests with a significance level = 0.05 (Figure 4B). Large mean differences with selbal were observed in all datasets tested across all DiCoVarML signature types. We observed moderate differences with coda-lasso for the CDI, NAFLD, and the Crohn’s classification tasks. Notably, for the HIV classification task, we only note significant moderate differences in AUC for the signature obtained by DiCoVarML with the Hybrid approach. We observed modest to moderate differences compared to clr-lasso on the CDI/Crohn’s classification task and large differences in AUC on the NAFLD and HIV classification tasks for all DiCoVarML signatures.

**Figure 4.**
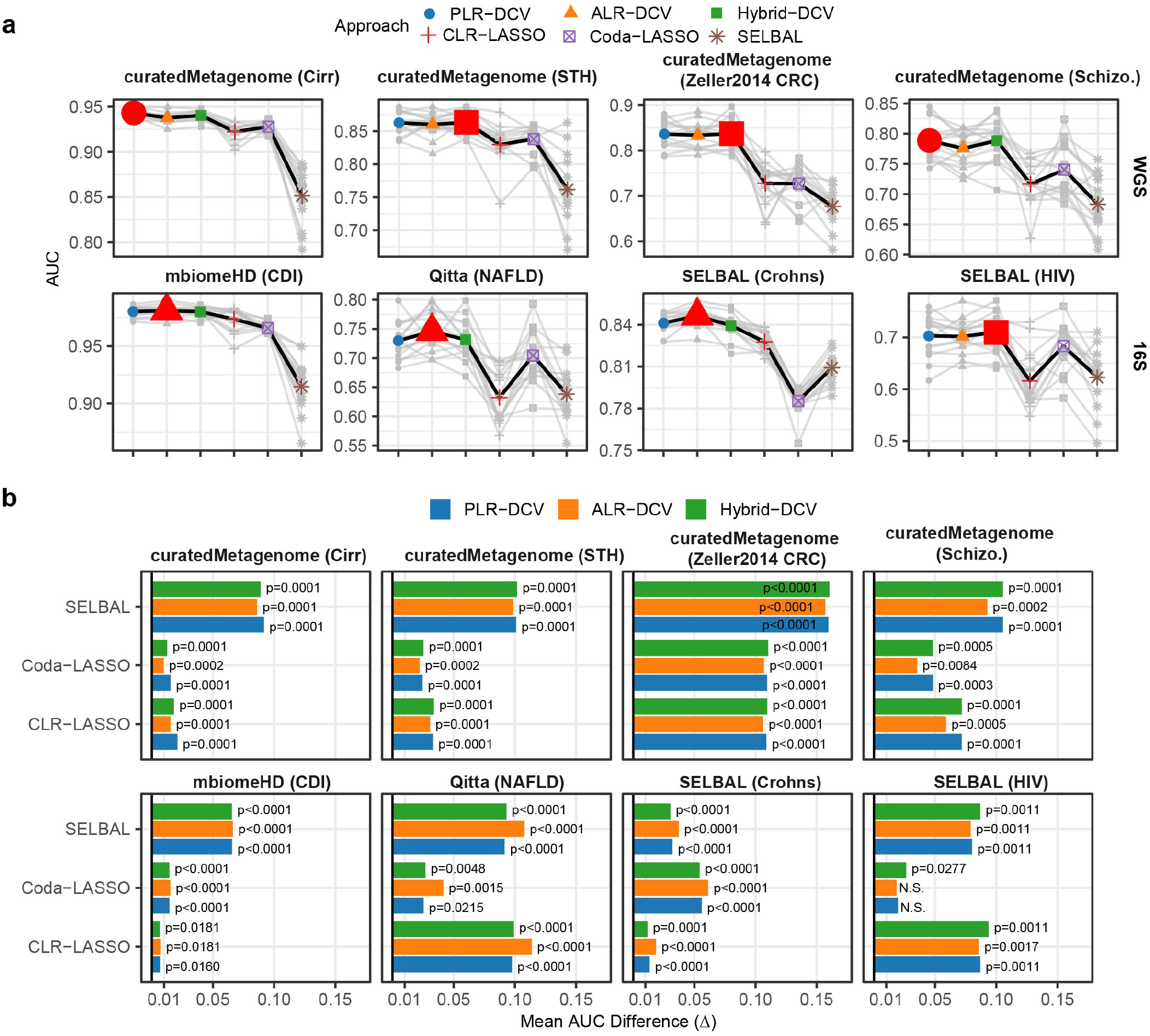
Classification performance comparison using real 16S and WGS datasets. Paired 15 repeats of 2-fold cross validation for each approach when applied to publicly-available case (shown) vs. control datasets. (a) AUC results for each method using WGS (top row) or 16S (bottom row) datasets. Grey points represent seed specific results. Grey lines connect paired seed specific AUC scores. Heavyweight points in color indicate overall mean AUC for each approach. Red point indicates approach with highest mean AUC. (b) Comparison of mean AUC differences between existing compositional approaches (y-axis) and DiCoVarML (bars). Results from Wilcoxon signed-rank test comparing AUC scores are shown. Benjamini–Hochberg-corrected *p*-values rounded to the nearest 0.0001 indicate high levels of significance in most cases (“N.S.” indicates not significant at the 0.05 level).

For WGS datasets, DiCoVarML signatures again achieved the highest mean AUC in all four datasets tested (Figure 4B), with PLR (2 of 4) and Hybrid (2 of 4) signatures achieving the highest mean AUC. Examination of mean differences compared to selbal revealed significant large mean differences in AUC across all datasets tested. For both coda-lasso and clr-lasso we observed significant yet modest to moderate differences for the cirrhosis (Cirr) and soil-transmitted helminth (STH) classification tasks and large significant differences for the Colorectal Cancer (CRC) and Schizophrenia (SCZ) tasks. Overall, our findings from real datasets demonstrate DiCoVarML signatures significantly outperform existing compositional methods for classifying disease using metagenomic data. In addition to achieving better classification performance, DiCoVarML signatures are sparse and easily interpretable. Furthermore, cross-validated performance estimates obtained from models trained with ALR or PLR are more robust and better representative of out-of-sample performance.

### DiCoVarML predicts onset of NEC in preterm-infants

In this case study, we apply untargeted DiCoVarML with PLR and ridge regression to predict the onset (> 7days) of necrotizing enterocolitis (NEC) in pre-term infants, using publicly-available fecal microbiome data from MicrobiomeDB^4^ collected as part of the longitudinal NICU NEC study^18^ (Figure 5). After data preprocessing (see Methods), the dataset contained 136 genera (384 unique taxa) across 902 (non-NEC = 779, NEC = 123) samples from 144 pre-term infants (non-NEC = 120, NEC = 24). First, we set out to understand if the microbiome compositions were predictive of future NEC. In order to unbiasedly evaluate the performance of classifiers in this imbalanced setting, we focus on AUC to assess performance across thresholds in a manner that is not biased to either the minority (NEC) or majority (nonNEC) group.^28^ We estimated the classification performance with 20 repeats of 5-fold stratified (by NEC status and sample) cross-validation using DiCoVarML with ridge regression to obtain a mean AUC = 0.676 (95% CI: 0.655 – 0.696) (Figure 5A). To ensure that patterns learned by the classifier were non-random, we computed the AUC under permuted disease labels within the same folds, achieving a mean AUC = 0.616 (95% CI: 0.595 - 0.636). Using the Wilcoxon signed rank test, we confirmed the true classifier performed better than random (*p* = 2.465 · 10^−5^) indicating the classification model learned non-random disease specific association patterns (Figure 5B/5C). To further explore these patterns and identify a candidate set of biomarkers, we trained the final DiCoVarML guided ridge regression on the full dataset, uncovering a microbial network connecting 8 genera by 14 ratios that is predictive of future NEC (Figure 5B), revealing increased abundance of *Staphylococcus, Klebsiella, Cutibacterium*, and *Gemella* relative to this microbial network were associated with future NEC onset. Using the final trained model, we next examined the regression scores for each sample relative to the number of days until NEC onset (Figure 5D), showing classification performance was strongest 11–17 days before onset with a notable decrease in accuracy 18 or more days before onset. Finally, analysis of survival agnostic regression scores from the NEC positive samples revealed a significant association (*p* = 4.566 · 10^−5^) between the regression score and survival with higher scores associated with samples from infants that did not survive (Figure 5E).

**Figure 5.**
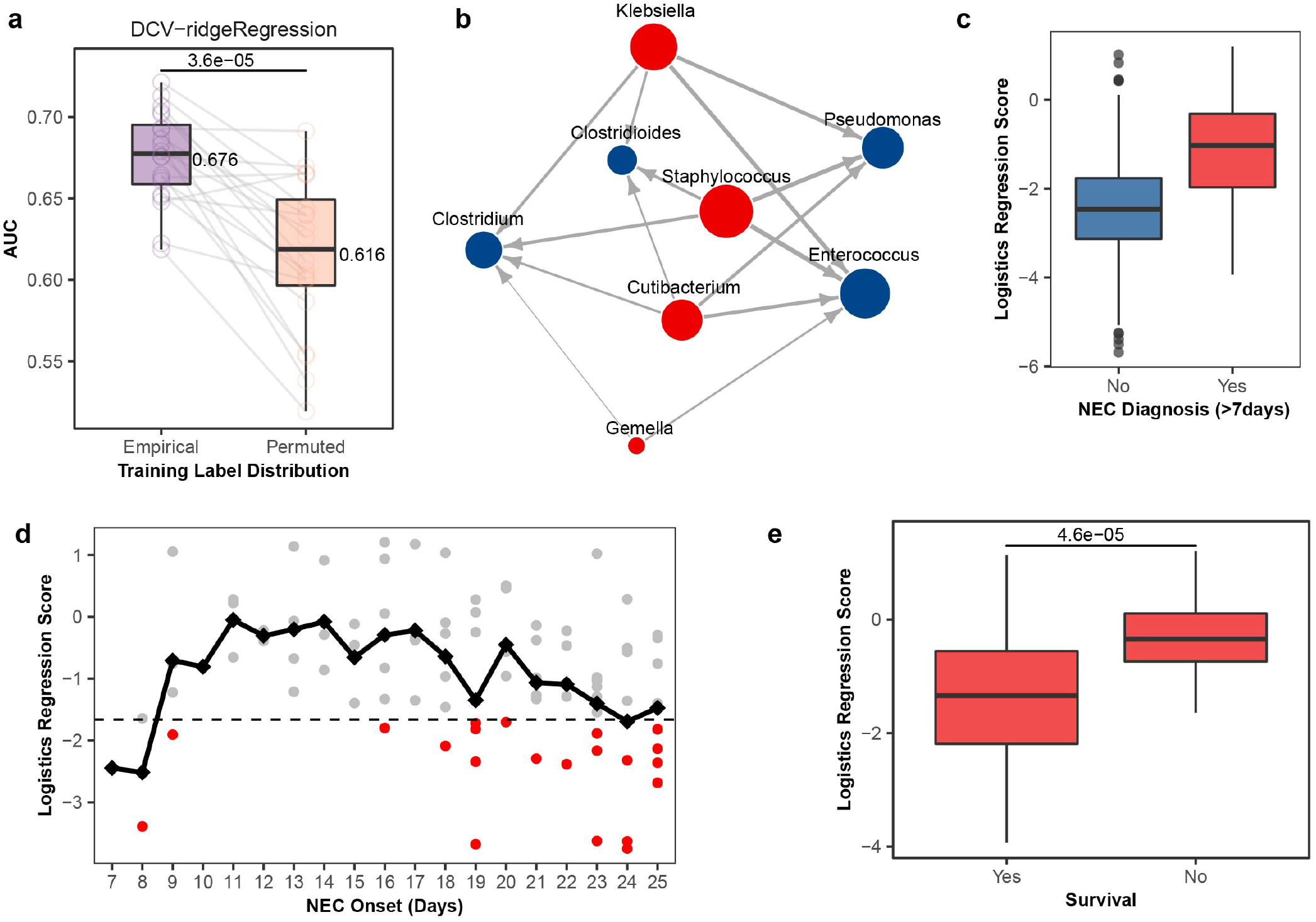
Predicting NEC onset (> 7 days) in premature infants using gut microbiome composition in the first few months of life using untargeted DiCoVarML with ridge regression applied to metagenomic profiled fecal samples (n=1,100) from the NICU NEC study. (a) AUC (paired by seed) from stratified (by sample and class) cross-validation (20 repeats x 5-fold) with empirical and permuted labels with *p*-value from a Wilcoxon signed-rank test and overall mean AUC shown. (b) Predictive microbial log ratio network after applying DiCoVarML to the full data. Nodes (v = 8) represent selected genera with sizes indicating the sum of absolute coefficients (*β*_*i*_) associated with each node. Each directed edge (e = 14) indicates a ratio (outgoing part over receiving part) with thickness proportional to the absolute coefficient *β*_*i*_ (larger absolute coefficient = thicker edges). Blue/Red node colors indicate increased abundance (relative to network) is associated with nonNEC/NEC samples. (c) Distribution of logistic regression scores 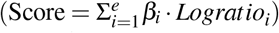 from model fit to full data. (d) Black line represents overall mean score by day. Dashed line indicates the non-NEC (lower) vs. NEC (above) decision threshold. Red points indicate misclassification (nonNEC vs. NEC). (e) Logistic regression scores of NEC positive samples only stratified by survival, with *p*-value from Mann-Whitney U test.

### Meta-Analysis with DiCoVarML reveals association between CRC, gut microbiome, and tumor staging

Lastly, we demonstrate the targeted DiCoVarML with ensemble modeling approach in a novel 11-cohort meta-analysis to classify fecal microbiome samples as colorectal cancer (CRC) vs. control. Specifically, we demonstrate how targeted (T = 50) DiCoVarML can be used to identify a predictive PLR signature (between species). Using the curatedMetagemic R package,^5^ we compiled and processed case-control WGS data from 11 studies (see Methods): keeping only samples with at least 10^6^ total reads, our final dataset included 1,305 samples (CRC = 653, control = 652). To estimate CRC vs. control classification performance we used 252 repeats of stratified (by dataset) cross-validation where, for each partition, both splits (5 and 6 of the 11 total datasets) were used once for training and testing. From this, we observed a mean AUC = 0.795 (95% CI: 0.791 – 0.798) which was significantly (*p <* 2.22 · 10^−16^) higher than the mean AUC = 0.498 (95% CI: 0.493 – 0.503) obtained with permuted labels (Figure 6A), confirming non-random predictive patterns. We next used DiCoVarML to infer, from the full dataset, a targeted 50 species PLR signature predictive of CRC. In doing this, we identified 259 log ratios between the targeted 50 species as being important for classification (Figure 6B). To understand the model predictions, we computed ensemble model scores (see Methods), revealing higher scores among CRC samples with control samples generally having lower scores (Figure 6C). After simplifying the log-ratio network, we next examined the multivariate association of individual taxa to each group (CRC/control) (Figure 6D), revealing *Peptostreptococcus stomatis, Gemella morbillorum*, and *Fusobacterium nucleatum* as the top three taxa associated with CRC, with increased abundance of these species within the microbial network (Figure 6B) associated with higher ensemble model scores (Figure 6C). Likewise, we found *Alistepes inops, Solobacterium moorei* and *Eubacterium eligens* to be the top three species associated with control samples (Figure 6D) where increased abundances of these species within the microbial network are associated with lower ensemble model scores. Finally, we tested if there was an association between the ensemble scores and cancer stage via AJCC (*n* = 258) or TNM (*n* = 168) staging. Indeed, using the stage-agnostic ensemble scores, we found strong associations between the ensemble score and stage for both AJCC (*p <* 3.696 · 10^−5^; Figure 6E) and TNM (*p <* 0.043; Figure 6F) labeled samples. Combined, our results indicate higher ensemble model scores are associated with advanced cancer stages, highlighting potential clinical utility of DiCoVarML.

**Figure 6.**
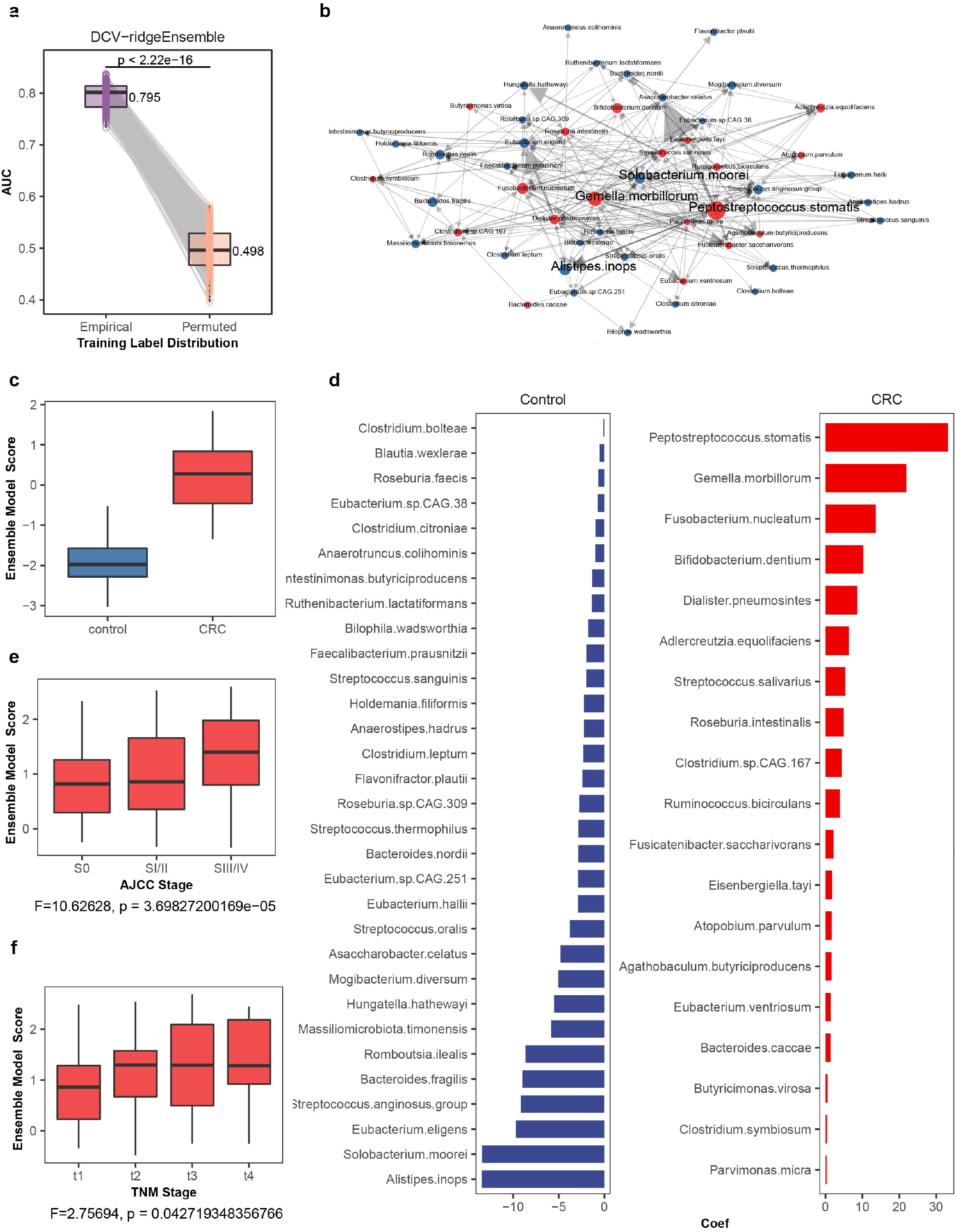
Meta-analysis classifying CRC vs. control from gut microbiome composition using targeted (T=50) DiCoVarML with ensemble model using samples (n=1,305) from 11 publicly-available cohorts. (a) AUC (paired by seed) with stratified (by dataset) cross-validation (252 splits x 2-fold) with empirical and permuted labels with *p*-value from a Wilcoxon signed-rank test and overall mean AUC shown. (b) Predictive microbial log ratio network after applying DiCoVarML to the full data. Nodes (v = 50) represent selected genera with sizes indicating the sum of absolute coefficients (*β*_*i*_) associated with each node. Each directed edge (e = 259) indicates a ratio (outgoing part over receiving part) with thickness proportional to the absolute coefficient *β*_*i*_ (larger absolute coefficient = thicker edges). Blue/Red node colors indicate increased abundance (relative to network) is associated with nonCRC/CRC samples. (c) Distribution of ensemble model scores from model fit to full data. (d) Species level contributions to ensemble model scores. Coefficient size (x-axis) indicates overall contribution to score where absence of nonCRC (blue bars) species are associated with CRC and the absence of CRC (red bars) genera are associated with nonCRC samples. (e) Boxplot of ensemble model scores stratified by AJCC stage (n = 258). (f) Boxplot of the ensemble model scores stratified by TNM stage (n = 168). *F*-statistics and *p*-values from Score ∼ Stage linear model.

## Discussion

In this paper we introduced and benchmarked the DiCoVarML framework for robust feature selection and classification of compositional datasets using log-ratio-transformed metagenomic features. Through our simulated example data scenarios and real-world data case studies, we have demonstrated the appropriateness and utility of this framework for supervised classification, as well as its particular relevance for metagenomic data. Importantly, our framework flexibly supports both targeted and untargeted feature selection in a multi-level manner to identify selected parts (e.g., taxa, metabolites) and a subset of log ratios between those parts. All of the code used to generate our results are included as part of the DiCoVarML R package developed to implement this framework.

## Methods

### DiCoVarML framework

We here describe the details of the various components of the DiCoVarML framework.

#### Data processing: sparse features and treatment of zeroes

The DiCoVarML framework takes as input taxonomic count tables from 16S or WGS sequencing, with or without zeroes. For taxonomic tables, reads should be assigned a taxonomy using a suitable database (e.g. SILVA, Greengenes, RDP, NCBI) and aggregated to a taxonomic level of interest (e.g. Genus, Family). DiCoVarML can handle any relative data where log ratio based predictors are of interest. In building models in terms of log ratios, one must necessarily select a strategy for handling zeroes and sparsely-counted taxonomic parts.

Sparse features are removed with a default 10% threshold, retaining taxa (parts) if present (counts ≥ 1) in at least 10% of samples. After sparse features have been removed, any samples with zero total reads on the remaining parts are also removed. While flexible, by default the DiCoVarML framework handles zeroes via the multiplicative replacement strategy described in Ref. 29 as implemented in Ref. 30 for metagenomic data. Importantly, because this strategy reinterprets a zero to mean that taxa are present but below a detection limit (uniform across parts), the log ratios between non-zero parts remain preserved. Moreover, in this strategy the log ratio between two zero-replaced parts becomes zero and thus contributes a zero value to a regression formula. Notably, while we find this strategy to be sufficient for robust predictions in our study here, zeroes can instead be imputed using others techniques such as those found in the zCompositions R package if desired.

In the application of our framework here, we aim to most conservatively minimize the possibility of any information leaking from test/validation sets leaking into discovery/train sets. To this end, we perform the removal of sparse parts/samples and the multiplicative zero replacement on both the training and discovery sets (described below).

#### Outer-loop cross-validation: classification performance

Because feature selection is an important step in the DiCoVarML framework, our cross-validation schema must account for this to minimize over-fitting. DiCoVarML uses nested feature selection to minimize the chance of reporting artificially high AUC estimates that might be obtained when feature selection is performed on the full dataset before cross-validation. Specifically, in frameworks where feature selection is not nested, inflation of AUC or other classification performance metrics can be directly attributable to information leakage from the test set during model training. The nested schema used in the DiCoVarML framework directly prevents such information leakage.

We estimate classification performance through *r* repeats of *k*-fold cross validation, stratified by group labels. Importantly, selection of the model type and number of parts within the DiCoVarML framework is nested and treated as a hyper-parameter that is tuned, with part and ratio feature selection performed as part of the training. Each fold is left out once for testing with the remaining folds used for discovery in the inner loop (described below). This process creates *k* unique discovery-test partitions for each repeat. For concreteness here, we denote the *i*th testing fold of the *j*th repeat as **ϒ**_*i j*_ and the union of the remaining folds in the *j*th repeat as the discovery set **Ψ**_*i j*_ =∪_*l* ≠*i*_ **ϒ**_*l j*_.

Each discovery set is further partitioned in the inner loop (as described below) to perform model and part selection. After fitting this model and selected parts to **Ψ**_*i j*_ (and re-selecting a subset of log ratios from the available parts as part of the fitting procedure), we then assess performance on the corresponding test set, **ϒ**_*i j*_. The overall classification performance estimate is obtained by averaging over all *i j* test fold indices. We focus here on AUC as the primary metric of interest, but other measures could also be used (e.g., AUPRC, accuracy, sensitivity, etc.).

We note that the computational costs involved in the DiCoVarML framework are highly dependent on the number of parts analyzed; reasonable computational time and memory requirements can be expected for *<* 500 parts after preprocessing.

#### Inner-loop cross-validation: model and part selection

Model selection, including selection of either a targeted or untargeted number of parts selection, occurs within an inner cross-validation loop. In this inner loop, we further partition the (outer) discovery set **Ψ**_*i j*_ into 2 folds, 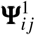 and 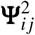, to be used as training-validation pairs. While this process can be repeated, the default DiCoVarML setting is to use a single repeat in this step to reduce computational time. Training on 1 of these 2 folds at a time, we identify a subset of *T* parts (e.g., taxa, metabolites, etc.) which then restrict the set of available log ratio features for model building. In our *targeted* feature selection mode, the parts subset size *T* is user defined. In contrast, *T* is selected automatically in the *untargeted* mode.

#### Targeted multi-level feature selection

The targeted multi-level feature selection method takes as input raw count data from a training set 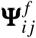, discovery set **Ψ**_*i j*_, or the full data for final model development (after out-of-sample performance has been estimated from the outer-loop cross-validation). Letting *p* be the number of parts after preprocessing, either 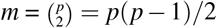 pairwise log ratios are computed for PLR signatures or the *m* = *p* − 1 additive log ratios are computed for ALR signatures forming the base log-ratio matrix **R**. For ALR/Hybrid signatures the reference part is selected such that given this reference, the Procrustes correlation between the ALR and the full PLR geometry is maximized.^15^ Part-level feature selection starts by computing the differential compositional variation (DCV) scores^30^ for each log ratio in **R**. The DCV scores are given by 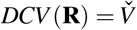. Specifically, DCV scores are computed using the dcvScores function from the selEnergyPermR R package.^30^ We then construct a weighted DCV network defined as 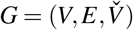 where *V* is the set of *p* part vertices, *E* is the set of *m* edges or pairwise log ratios between parts, and 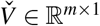 are the weights given by log ratio DCV scores. Let the (unweighted) adjacency matrix **A** ∈ ℝ^*p*×*p*^ and weight matrix **W** ∈ ℝ^*p*×*p*^ be derived from *G*. We compute the vertex strength for the *u*th vertex (representing the *u*th part) as 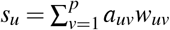 We select the top *t*_*i*_ parts from these strengths and compute all 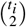 pairwise log ratios between them for PLR/Hybrid signatures or all *t*_*i*_ − 1 additive log ratios (with a Procrustes correlation maximizing reference) for ALR signatures to form the targeted log ratio matrix **L**. Log-ratio-level feature selection is then carried out by first computing the DCV scores for each log ratio, retaining only those with positive DCV scores. With this, the subset of log ratios is selected by sequential addition from the list of log ratios sorted by DCV score (high DCV scores first) until the subset includes at least *t*_*i*_ parts.

#### Model and target part selection

Using inner-loop cross-validation the best model and number of target parts can be selected by averaging the AUC estimates across the {train, validate} sets. From these results we select the model (*m*_*max*_) and number of target parts (*t*_*max*_) with the highest average AUC to to estimate classification performance (across discovery sets **Ψ**_*i j*_) or build a final model (using full data).

#### Machine learning models

While the choices of which machine learning model paradigms to use within DiCoVarML is flexible and can be determined by the user, we have here focused on the ridge regression and ensemble model modeling paradigms as the defaults to be utilized in DiCoVarML. Ridge regularized logistic regression in DiCOVarML uses the glmnet R package with alpha = 0 and type set to either binomial for binary classification or multinomial for multi-classification. Cross-validated AUC values are computed and stored. For final model development, regression scores and coefficients can be directly interpreted from the model using either the predict() or coef() functions from the R STATS package.

For the average probability ensemble model that we use by default in DiCoVarML, we apply the following machine learning algorithms: random forest (ranger R package), support vector machine with radial basis function (kernlab R package), regularized regression (glmnet R package), random forest with extra trees (ranger R package), and partial least squares discriminate analysis (pls R package). Each model is fit (to the appropriate fold) and the predicted class probability 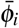 for the *i*th sample (e.g., in application to the corresponding validation or test fold) is computed as the average probability across all models used. Cross-validated AUC values are computed and stored. For all machine learning modeling the Caret R package is used. All models are trained with default parameters except random forest models are trained with 500 trees and mtry tuning grid using either 1 or the square-root of the number of log ratio features.

#### Final model scores and log ratio coefficients

In DiCoVarML, final model scores are computed after cross-validation performance has been estimated. For ridge-regression models, the final model score for each sample is obtained after fitting a ridge regularized logistic model to all data (after processing to remove sparse features/samples and zeros) using the cv.glmnet() from the glmnet R package with alpha = 0 and default settings otherwise. The fitted *β* coefficients correspond to log ratio features selected for the final log ratio signature.

Ensemble model scores for binary classification are obtained after fitting the ensemble model (see above) to all data (after processing). While direct interpretations of ensemble models are difficult, in DiCoVarML we transform each average prediction probability 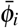 to an unconstrained ensemble model score 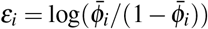, imputing non-zero values in the numerators or denominators of this transformation by the same procedure described previously for zeros in the data. We then fit a secondary ridge penalized linear regression model (adjusting for class/group) to understand how each log ratio in the data contributes to *ε*, using the glmnet R package with default *λ* grid. The resulting model coefficients may be used to interpret how each log ratio contributes to the final ensemble model score.

### Synthetic compositional data generation

In this section we describe in detail how we generate synthetic data and decision boundaries for the examples in Figure 2 and the synthetic data distribution benchmarks in Figure 3. Notably, decision boundaries can be described as hypersurfaces that separate a space into different classes/groups. We generate binary classification problems in our synthetic examples and benchmarks. We also describe the machine learning analysis used to assess classification performance.

#### Isoproportional decision boundary

Given a three part composition (*p, q, r*), an isoproportional line representing a fixed ratio between 2 parts is a straight line on the 2-dimensional simplex. That is, the isoproportional line between 2 parts (*p, r*) is defined by the relationship *p/r* = *C* for constant *C* (independent of *q*). We simulate two-class compositional data from a simple additive logistic normal distribution^8^ with *D* = 3 parts (*p, q, r*) as follows. Working in the *d* = *D* − 1 = 2 dimensional space of log ratios 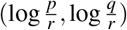, we define the mean vectors (**m**_1_, **m**_2_) ∈ ℝ^*d*^ and covariance structures (**Σ**_1_, **Σ**_2_) ∈ ℝ^*d*×*d*^ for each class. The simulated training data in the top row of Figure 2 was generated from multivariate Gaussian distributions with means **m**_1_ = (−0.5, 2.0) and **m**_2_ = (0.5, 2.0) and covariances diag(**Σ**_1_, **Σ**_2_) = 0.2 and off-diagonal covariance elements drawn uniformly at random on [0.0, 0.1], then ensuring the covariance matrices are semi-positive definite by application of the nearPD() function from the R Matrix package. We note in particular that these simulated points correspond to a ground-truth isoproportional boundary *p/r* = 1 (i.e., 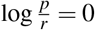). Finally, we apply the one-to-one additive logistic transformation to map each simulated log ratio point ***r*** ∈ ℝ^*d*^ back to ***x*** ∈ ℝ^*D*^ in the space of compositions by *x*_*i*_ = exp(*r*_*i*_)*/S* for *i* = 1, …, *d* and *x*_*D*_ = 1*/S*, with normalization constant *S* = exp(*r*_1_) + · · · + exp(*r*_*d*_) + 1. The test data in the top row of Figure 2 was generated by the same process as the training data but with means **v**_1_ = (−0.5, −2.0) and **v**_2_ = (0.5, −2.0). Importantly, the positions of the test means correspond to the same isoproportional boundary as the training data but centered in a different region of the simplex due to different-scaled contributions from the *q* component, representing, e.g., possible out-of-study differences across populations, methodologies, or noise levels.

#### Compositional line decision boundary

Using the so-called “Aitchison’s Geometry,” compositional boundaries on the simplex can be defined using geometric power transformations and perturbation operations.^31^ Given a compositional vector **z** = [*z*_1_, …, *z*_*D*_] on a (*D* − 1)-dimensional simplex (that is, constrained to ∑_*i*_ *z*_*i*_ = 1), the power operation is defined by *α* 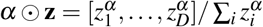. Similarly, the perturbation operation between two compositional vectors **y** and **z** is defined by **y** ⊕ **z** = [*y*_1_*z*_1_, …, *y*_*D*_*z*_*D*_]*/* ∑_*i*_ *y*_*i*_*z*_*i*_. Armed with these algebraic operations, we can define a compositional-line decision boundary **b**(*α*) = **y** ⊕ (*α* ⊙ **z**) by specifying **y** and **z**, with *α* ∈ [−∞, ∞]. The boundary in the bottom row of Figure 2 was defined by setting **y** = (1*/*3, 1*/*3, 1*/*3) and **z** = (0.6, 0.3, 0.1).

For interpretation, we emphasize that a decision boundary constructed in this way is equivalent to the zero level set of particular linear combinations of log ratios, noting that different linear combinations of log ratios may be algebraically equivalent to one another. Indeed, such linear combinations of log ratios are then equivalent to a linear combination of log part values which are automatically constrained to be independent of a constant multiplicative factor applied uniformly to all values. In particular, in terms of the composition (*p, q, r*), the decision boundary in the bottom row of Figure 2 using the **y** and **z** listed above is equivalent to the zero level set of 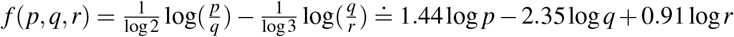. That is, *f >* 0 is on one side of the boundary with *f <* 0 is on the other side. We again note there are other linear combinations of log ratios giving the same zero level set boundary. We call attention to the fact that the resulting coefficients on the logged parts (i.e., on log *p*, log *q*, log *r*) sum to zero by construction, since any linear combination of log ratios is independent of an overall multiplicative scale applied uniformly to each component part. We also note that the decision boundary in the bottom row of Figure 2 has zero offset because of the symmetry in the selected **y** used here, but non-zero offsets would result from other **y** selections. Finally, we note that the zero level set of a linear combination of log ratios (with possible non-zero offset coefficient) is left unchanged by a scalar multiple applied across all coefficients; that is, in a sense aligned with logistic regression, the *f* = 0 boundary of equal probability would be left unchanged but the certainty one might assign to a class label prediction is affected by how quickly *f* changes away from the zero level set.

To generate simulated data (perfectly) separated by the compositional line boundary **b**(*α*), we generate *n* = 100 random *α* values uniformly on [−10, 10] and evaluate **b**(*α*) for each point. We then randomly shift each of these points to the same side of the boundary by sampling the shift size Δ uniformly on [0.1, 0.2] and moving the point to **x** = **b**(*α*) ⊕(Δ⊙ ***ρ***), where the constant shift direction here is set to ***ρ*** = (1, 1, 100)*/*102. We similarly generate points on the other side of the boundary by the same procedure, starting from *n* = 100 new *α* values and a new Δ for each point, but with the shift given by **x** = **b**(*α*) ⊕ ([−Δ] ⊙ ***ρ***). Finally, we partition this data into train and test sets according to the value of the first component (*x*_1_ = *p* in (*p, q, r*)) of each data point, with *x*_1_ *>* 1*/*3 points assigned to the train set and those with *x*_1_ ≤ 1*/*3 assigned to the test set.

#### Machine learning analysis

For the isoportional decision boundary simulation 100 samples were generated for both classes of data for both training and testing sets. Using the R caret and Random Forest packages, random forest models were fit to the training data with mtry = 2 with ntrees = 500. Training set AUC was estimated using 10 repeats of 10-fold cross-validation. Final models for all data transformations were fit to the full training dataset and used for prediction on the testing dataset. Classification performance was estimated using the AUC metric from the R pROC package.

For the compositional line decision boundary, 200 samples for each class were simulated for both training and testing sets. Using the training set, feature discovery and model fitting was carried out with the glmnet R package with alpha = 1 (LASSO penalty). Only the selected features from these procedures were included in the final training and test data. Notably, to simulate true feature selection, such as in the case of biomarker discovery, for relative abundance normalized and CLR-transformed data, the normalization/transformation operation was applied to the selected features. For PLR-transformed data, only the selected ratios were computed on the training and test data. A final logistic regression model was fit to the training data and tested on the test data for all data transformations. Classification performance was assessed using the AUC performance metric.

### Synthetic 16S and WGS data simulation

To simulate data with real 16S and WGS characteristics, we used the selEnergyPerm R package. Specifically, we used the simFromExpData.largeMeanShft() function from the selEnergyPermR package to simulate count data with differences in the mean vectors of additive log ratio transformed data. To do this, we used the process described in Ref. 30 where a zero inflated negative binomial model (ZINB) is fit to healthy WGS sequenced samples from the Observational Study of Blood Glucose Levels and Gut Microbiota in Healthy Individual trial^32^ using the curatedMetagenomicData^5^ and 16S sequenced fecal samples from the TwinsUK population^33^ using the zinbwave R package. We note parametric simulations and goodness of fit analysis in Ref. 34 demonstrate ZINB models are reasonable for modeling metagenomic count data. Once the ZINB models were fit we synthesized binary class count data for *n* = 200 samples (100 in each class), with true differences between the classes. In addition to the raw count matrix, the primary experimental parameters of the simFromExpData.largeMeanShft() function are the featureShiftPercent parameter (effect size) which controls the magnitude of difference between mean vectors of each class and the perFixedFeatures parameter (Signal Density) which controls the number of parts that will have simulated between class mean differences where higher values are associated with fewer parts being shifted (sparse signal density). From a machine learning classification perspective, these parameters directly control how difficult or easy the learning task will be where smaller effect sizes among fewer parts are more difficult to learn vs. larger effect sizes among more parts which are easier to detect. Here we set featureShiftPercent = {18%, 24%, 30%} and perFixedFeatures = {50%, 70%, 90%, 95%, 98%}.

### Publicly-available datasets used for benchmarking

We utilized 16S and WGS data from several publicly-available data sources as benchmark data, extracting processed OTU/taxa tables with matched clinical data. The following data sources were used:

#### curatedMetagenomicData

Metagenomic datasets were extracted from version 3.0.0 of the curatedMetagenomicData^5^ R package. The curatedMetage-nomicData R package provides a common interface to access standardized metagenomic data (e.g. relative abundance, pathway abundance) for new analyses. Using the curatedMetagenomicData() function in this package, the following studies were extracted by study ID: ZhuF_2020,^35^ RubelMA_2020,^36^ WirbelJ_2018,^22^ and QinN_2014.^10^ The SummarizedExperiment R package rowData() and assay() functions were used to extract the taxa information and count tables respectively. Sample clinical data were extracted directly from the curatedMetagenomicData object using the SummarizedExperiment colData() function. Each study was checked for repeated measures on subjects to ensure appropriate cross-validation schemes were applied. Only samples with at least 10^6^ total reads were included in our analysis. Additionally, taxa were aggregated to the species taxonomic level.

#### Qitta

Metagenomic data for the gut microbial metabolism and nonalcoholic fatty liver disease (NFALD) study^37^ were downloaded from Qitta.^6^ Qitta is an open source multi-omics platform that provides database resources for storing and processing publicly-available omics studies. Using this platform, the sample information and two related BIOM files were downloaded directly from the web interface using Qitta Study ID 11635. The 16S BIOM files were loaded in R using the biomformat R package. For this study, the total samples (*n* = 290) were filtered to only include samples with a NFALD status collected with at least 1,000 total reads, yielding 185 samples (NFALD+Cirrhosis = 44, nonNFALD+Cirrhosis=141). Taxa were aggregated to the genus taxonomic level.

#### MicrobiomeHD

16S data from the clostridium difficile infection (CDI) microbiome study^38^ were downloaded from the microbiomeHD database,^39^ a standardized database of uniformly processed publicly-available case-control studies. Specifically, the *cdi_schubert_results*.*tar*.*gz* file was downloaded from the microbiomeHD web interface. Within this directory, we extracted the OTU data from *cdi_schubert*.*otu_table*.*100*.*denovo*.*rdp_assigned* and sample clinical data from *cdi_schubert*.*metadata*.*txt*. Taxa data were aggregated up to the genus level.

#### Selbal

The 16S Crohn’s and HIV benchmark datassets originally included in the presentation of the selbal method^16^ were extracted from the cloned selbal R package available on gitHub.^16^

### Publicly-available case study datasets

#### Necrotizing enterocolitis (NEC)

Fecal metagenomic data were extracted from microbiomeDB version 23,^4^ a web based discovery tool and database that uniformly processes and stores 16S and WGS datasets along with related clinical metadata, using the NICU NEC (WGS) study ID. The longitudinal NICU NEC study^18^ contains 1,100 microbiome-profiled fecal samples from 150 pre-term infants collected during the first months of life where some infants go on to develop NEC. For our case study, we downloaded processed WGS taxon abundance (*NICUNEC*.*WGS*.*taxon_abundance*.*tsv*) and sample detail (*NICUNEC*.*WGS*.*sample_details*.*tsv*) tables. The NEC positive samples were filtered to include only microbiomes related to an onset of greater than 7 days (Days of period NEC diagnosed > 7) and samples after the first day (Age at sample collection days > 0) of life (since some samples developed NEC within the first 7 days of life). The filtered NEC positive samples were then merged with the NEC negative samples. The taxa tables were aggregated at the genus level. Further, because these data are longitudinal and involve repeated measures, cross-validation folds were evenly stratified by infant and NEC.

#### Colorectal cancer (CRC) meta-analysis

Fecal metagenomic datasets for our 11-cohort meta-analysis were processed and collected from the curatedMetagemic R package (also used for some of the benchmark datasets, as described above). In particular, we aggregated sample clinical data and WGS fecal microbiome data (*n* = 1, 395) from the following study IDs: FengQ2015 (*n* = 107),^19^ GuptaA2019 (*n* = 60),^20^ HanniganGD2017 (*n* = 55),^21^ WirbelJ2018 (*n* = 125),^22^ ThomasAM2018b (*n* = 60),^23^ ThomasAM2018c (*n* = 80),^23^ VogtmannE2016 (*n* = 104),^24^ ThomasAM2018a (*n* = 53),^23^ YachidaS2019 (*n* = 509),^25^ YuJ2015 (*n* = 128),^26^ ZellerG2014 (*n* = 114).^27^ Only samples with at least 10^6^ total reads were retained (*n* = 1, 305). To most conservatively estimate out-of-sample generalizability, all (outer loop) cross-validation partitions in this meta-analysis were strictly stratified by dataset only. Because only 9 samples from the HanniganGD2017 were retained, we always included this dataset in the partition with 6 data sets (the other partition containing 5 data sets). Our analysis then considered all 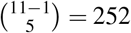 such partitions into 5/6 datasets, using each split once for training and testing.

### Use of existing methods in benchmarks

#### Selbal

To estimate the classification performance of selbal we used the selbal R pacakge available on github.^16^ Because the primary selbal function in the selbal package performs cross validation internally, the following process was used to comparably nest the selbal feature selection method and test on common folds. Using the count data from the *i j*th discovery set, we identify taxa to include in the balance using the selbal.aux() function with the zero.rep = “one”. If the number of taxa returned after this step is ≤ 1 then all taxa are included. After taxa have been identified we extract the balance values for both the discovery and testing folds using the bal.value() function. A final logistic regression model is fit to the balances with corresponding class labels for the discovery set and used to make predictions on the test. Performance metrics are computed as described above.

#### CLR-LASSO

To implement the clr-lasso method we use standard functions in a manner consistent with Ref. 17. To comparably apply the clr-lasso method in a nested way to common folds for benchmarking, we first compute the CLR transformation on the discovery/test sets after adding a pseudo count of 1. Feature scaling is first estimated from the discovery set and then applied to scale features in the discovery and test sets. The glmnet R package is then used to fit a LASSO model (alpha = 1) to the discovery data for feature discovery. If the number of features selected after this step is ≤ 1 (indicates LASSO failed to converge) then all features are used. After subsetting the discovery and test sets for the *i j*th parititon to include only the features selected (nested feature discovery), the CLR transformation is recomputed. Finally, after scaling features as described above, a LASSO penalized logistic regression model is fit to the discovery data (after feature selection) and used to make predictions (i.e. using the predict() function with “lambda.min”) on the test data (after feature selection). Overall performance metrics are computed as described above.

#### Coda-LASSO

To implement the coda-lasso method for feature selection we cloned functions from the referenced github repository in Ref. 17 and carried out the analysis similar to the github documentation. Specifically, to comparably estimate classification performance in a nested manner on common folds, the following process was used. We first added a 1 pseudo count to the discovery and testing data sets (default for coda-lasso). Here, to minimize test data leaking into the discovery set, the z-transform (coda-lasso specified) was estimated from the discovery data only and applied to both the discovery and test data for the *i j*th partition. To identify a suitable lambda value for input into the coda_logistic_lasso() function, we used the glmnet R package to estimate a grid of lambda values and then selected the lambda value that both included *>* 0 features and maximized the “proportion of explained deviance” estimated from the coda_logistic_lasso() function. Once lambda was estimated, the coda_logistic_lasso() was used to select a subset of features for further model development. After subsetting the discovery and test sets for the *i j*th partition to include only the features discovered in the previous step, a pseudo count of 1 was added and the z-transformation was again performed as described above. If the number of features selected after this step was ≤ 1 (indicates LASSO failed to converge) then all features were used. We again optimize lambda and construct a final LASSO regularized logistic regression model using the coda_logistic_lasso(). The returned “beta” coefficients from the coda_logistic_lasso() function were used to compute the final scores for both the discovery and test data. From this, a logistic regression model was fit to the discovery data (after feature selection) and used to make predictions on the test data. Overall performance metrics are computed as described above.

## Data availability

All data used to benchmark the DiCoVarML framework is publicly available and can be accessed as described in the Methods section.

## Code availability

All functions required to implement the DiCoVarML framework have been made publicly available in the DiCoVarML R package available on gitHub at https://github.com/andrew84830813/DiCoVarML.git. All code used here to benchmark the DiCoVarML framework, conduct case studies, and produce the figures is accessible on github at https://github.com/andrew84830813/DiCoVarML.ProjectRepo.git.

## Acknowledgements

We are grateful to our collaborators for many conversations that helped motivate and further refine the development of this framework, including Zachary Boyd, Wesley Burks, Jeff Henderson, Elise Hickman, Ilona Jaspers, Mike Kulis, Laura Marks, John Robinson and William Weir.

## Author contributions statement

ALH developed the framework methodology, performed simulations and case studies, developed the software package, and wrote the manuscript. PJM provided advice and feedback during the development of the framework methodology and wrote the manuscript. ALH and PJM approved the final manuscript.

